# The *panC*-encoded pantothenate synthetase to tackle carbapenem-resistant OprD *Pseudomonas aeruginosa* mutant revealed through Tn-Seq

**DOI:** 10.1101/2023.10.19.563089

**Authors:** Cléophée Van Maele, Ségolène Caboche, Marin Moutel, Arnaud Bonnomet, Sophie Moussalih, Emilie Luczka, Hervé Jacquier, Anaëlle Muggeo, Thomas Guillard

## Abstract

For the World Health Organization, carbapenem-resistant *Pseudomonas aeruginos*a is a critical priority for which new antimicrobial drugs are needed. Consequently, understanding the underlying mechanisms of resistant bacteria infection will enable the identification of new therapeutic targets. Loss of the OprD porin is the main determinant of resistance to the last resort carbapenem antibiotics and has been described to enhance fitness *in vivo* and virulence. Transposon sequencing is a high-throughput sequencing technique that makes it possible to identify essential genes (EGs) that may turn out to be therapeutic targets. However, such a strategy has not yet been used for OprD-deficient *P. aeruginosa*. In this study, we identified the EGs specific to PA14 OprD mutant for LB growth and we established a list of 30 EGs among these, we highlighted the *panC* gene encoding pantothenate synthetase as a promising target. Using CRISPRi, we confirmed that silencing *panC* reduced LB growth, and decreased *sigX* expression, whose overexpression is associated with membrane fluidity, as well as the expression of genes involved in the fatty acid synthesis (FAS). Taking into account the weakness of PA14 OprD mutant due to an altered membrane consecutive to a decrease in unsaturated FAS in the absence of *panC,* we showed that silencing *panC* extended the destruction time of 16HBE airway cells. Overall, our findings highlighted the anti-virulence potential of *panC* inhibition and shed new light on its inhibition as a target for treating carbapenem-resistant OprD-defective PA lung infections.

## Introduction

*Pseudomonas aeruginosa* (PA) is an opportunistic pathogen known to cause various types of infection. The main one is pulmonary infection, such as pneumonia when associated with respiratory pathologies such as cystic fibrosis (CF) or Chronic Obstructive Pulmonary Disease (COPD) (1–3). PA is often found in hospital settings, particularly intensive care units, which favors the presence of multidrug-resistant (MDR) bacteria (4), a growing problem due to the lack of therapeutic choices for treating these infections.

PA is intrinsically resistant to many antibiotics and can acquire resistance upon treatments. For this reason, PA is part of the ESKAPE pathogens (5). The carbapenems are major antibiotics used as last resort treatments to face MDR PA. The lack of functional OprD porin leads to carbapenem resistance and is the most frequent mechanism of resistance to this class of antibiotics (6–8). OprD is collaterally the entry channel for two carbapenems (Imipenem and Meropenem), its first function is the uptake of basic amino acids such as arginine, lysine, histidine and ornithine, or even small peptides (9,10).

Interestingly, an enhanced *in vivo* fitness of OprD PA mutant has been described in the context of gastrointestinal and spleen colonization (11,12). It has also been shown to be more virulent in a mouse model of acute pneumonia (13). However, the mechanism underlying this increased virulence in the absence of OprD has yet to be elucidated. We therefore decided to determine whether *in vitro* gene essentiality could provide clues to tackle such a higher virulent and resistant pathogen.

Transposon Sequencing (Tn-Seq) is a high-throughput sequencing that uses transposon mutant libraries to determine the fitness or essentiality of genes under specific conditions. The use of a mariner transposon that inserts into a thymine-adenine (TA) site only once per bacterium and randomly makes it possible to obtain a very large number of mutants (14). The determination of essential genes (EGs) can lead to the identification of potential new therapeutic targets.

In this study, we first propose to evaluate the bioinformatics analysis of Tn-Seq data with data generated on two strains growing on Lysogenic Broth (LB): PA14 wild-type (WT) and PA14Δ*oprD*. As the regularly-updated TRANSIT software, offering several statistical methods and parameters, is the most popular tool, we decided to evaluate several methods and parameters of the TRANSIT suite. We also evaluated FiTnEss (Finding Tn-Seq Essential genes), a tool developed for dealing with PA Tn-Seq data (15). We introduced a gold standard dataset of EGs for PA14 and used several metrics to compare the different methods. From these Tn-Seq analyses we identified *panC* as a specific essential gene for the OprD mutant. We showed that the lack of import by the OprD porin did not explain the need of *panC* for *in vitro* growth. Nevertheless, we found that *panC* inhibition impacted the fatty-acid pathways indicating a decrease in unsaturated FA synthesis, which and could therefore affect the membrane homeostasis. Eventually, we evidenced that silencing *panC* decreased the virulence of the carbapenem-resistant PA OprD mutant against 16HBE cells and could be a promising therapeutic target.

## Results

### Description and quality of the data

Sequencing was carried out in 2 different runs: samples S1 and S3 on RUN1, samples S1-S2 and S5-S6 on RUN2 leading to 6 samples corresponding to PA14 WT or PA14Δ*oprD* grown on LB culture media (Table 1). Strains information are listed in table S1.

**Table 1:** Description of samples used in this study.

| RUN | Samples | Strain | Culture medium |
| --- | --- | --- | --- |
| RUN1 | S1 | PA14Δ*oprD* | LB |
| RUN1 | S3 | PA14Δ*oprD* | LB |
| RUN2 | S1 | PA14 WT | LB |
| RUN2 | S2 | PA14 WT | LB |
| RUN2 | S5 | PA14Δ*oprD* | LB |
| RUN2 | S6 | PA14Δ*oprD* | LB |

We showed that the quality metrics obtained with BWA (mapper in TPP package of TRANSIT) (16) were very different, being below the recommended thresholds, from those observed with bowtie (17) (Supplementary Table S2 and S3). Although, there are no formal criteria for determining what constitutes a good library and for excluding certain samples (18), a very important metric is density, which should be above 30 % and ideally above 50 %. In our study, density was above 50% for all the samples. The percentage of mapped reads was lowest for sample S6 with around 65% of reads mapped. In addition, sample S6 showed NZ-mean values of 14.4, which ideally should be greater than 50. To identify potential outliers, a correlation matrix (Pearson coefficient) was computed (Supplementary Figure S1). The results showed that correlation coefficients were higher between the same strains. With regards to replicates, the coefficients were lower with sample S6, which was therefore discarded. Finally, we used samples S5_RUN2, S1_RUN1 and S3_RUN1 for the PA14Δ*oprD* LB condition and samples S1_RUN2 and S2_RUN2 for the PA14 WT LB condition.

### Bowtie_HMM and FiTnEss for reliable EGs identification of PA14 in LB

Many previous studies have reported the EGs for several PA strains grown on different culture media (Supplementary Table S4) (11,15,19–21). First, we attempted to provide the minimal set of EGs of PA14 grown in LB, being somehow the gold standard EGs, which are expected to be retrieved in any set of EGs identified. From previous studies, 115 genes (GOLD_115) were common to all studies for PA14 and by intersecting all datasets, we obtained 84 genes (GOLD_84) that represented a reduced but reliable list of EGs.

To determine which analysis will provide the most reliable set of EGs from our Tn-Seq data, we compared the finely tunable TRANSIT suite with FiTnEss. Based on our GOLD_115 and GOLD_84 lists, we compared the EGs obtained from samples S1 and S2 (PA14 WT) using bowtie and BWA with different parameters and normalization. First, for both gold standard lists of EGs, less than 10% of genes were retrieved with BWA using Gumbel and HMM methods whereas 40 to 50% were retrieved with bowtie using either Gumbel or HMM. (Fig. 1A). Interestingly, considering EG and “growth defect (GD) genes, the HMM method identified around 90% of gold standard EGs for both mappers (Fig. 1A). Second, using bowtie with Gumbel or HMM, varying several parameters (see Material and Methods as well as the Supplementary data), we confirmed that Gumbel led to shorter lists of EGs whatever the parameters (Fig. 1B and Fig. S3). Surprisingly, we observed that using the mean read counts with Gumbel increased results by 10%. For HMM, using the default mean read counts was better than using the sum. Using the loess option predicted slightly more genes (194 EG and 602 EG+GD with loess option against 184 and 565 without loess option, respectively) but had no impact on the percentage of gold standard EGs retrieved. Third, as normalization of the read counts prevented other sources of variability from being mistakenly treated as real differences in datasets, we tested the several types of normalization proposed by the TRANSIT suite (Totreads, TTR, Nzmean, Quantile, Betageom, Zinfnb and nonorm) for bowtie_HMM. We found the highest amount of (EG + GD) with Quantile (690 and 2815) and Zinfnb (901 and 2815) normalization (Fig. 1C), but the number of identified genes were higher and therefore probably contained more false positives (Fig. S4). The default TTR normalization preserved good sensitivity and specificity.

**Figure 1.**
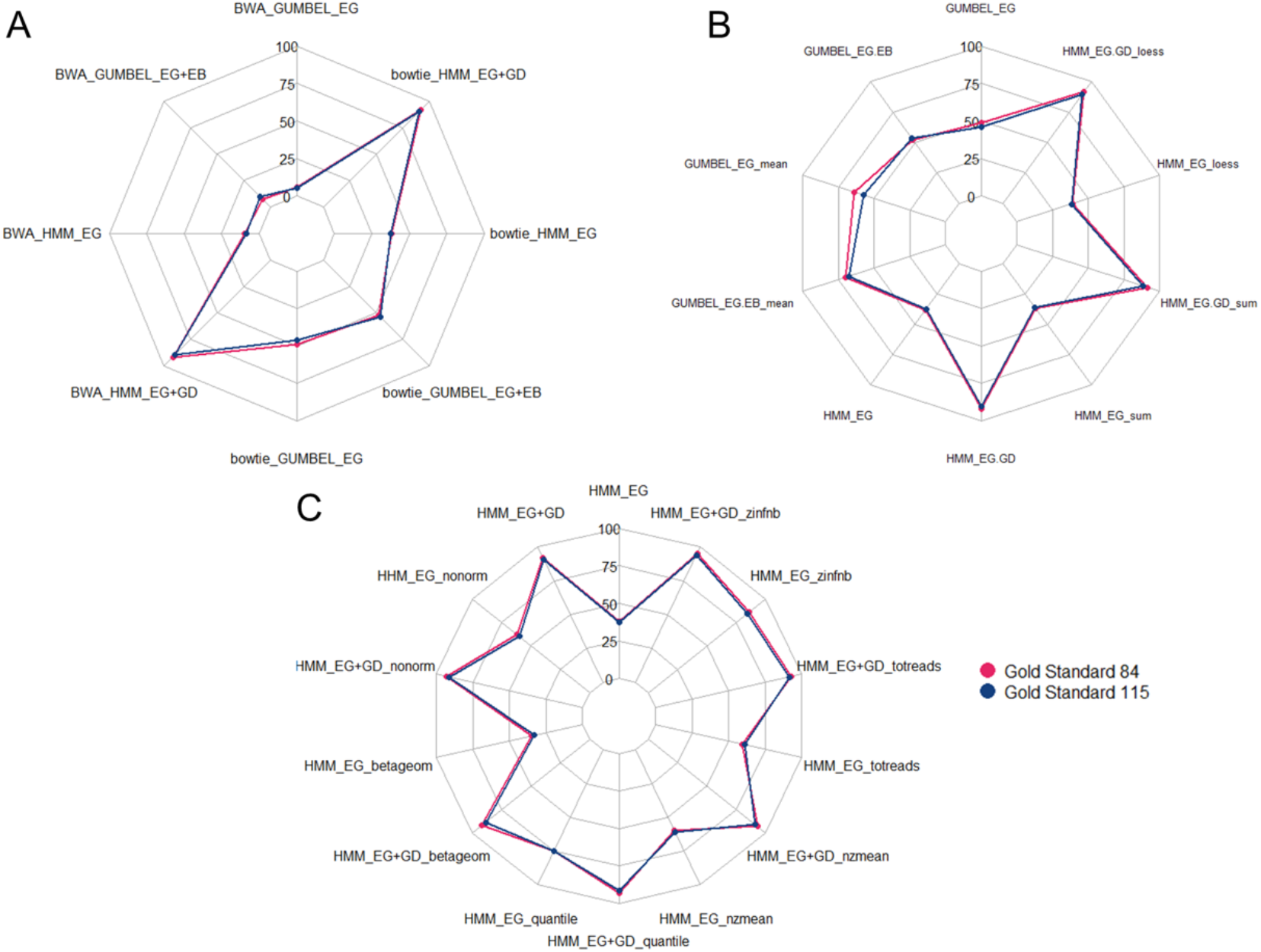
Assessment of Tn-Seq analysis methods. **(A)** Impact of the mapper on EGs determination illustrated by a radar plot showing the percentage of gold standard EGs identified by each method. **(B)** Impact of the different software parameters on EGs determination illustrated by a radar plot showing the percentage of gold standard EGs identified by each method **(C)** Impact of normalization on EGs determination illustrated by a radar plot the percentage of gold standard EGs identified by each method. For each panel: in blue the data set containing the GOLD_84 and in red the GOLD_115 from WT grown in LB. EG = Essential class, EB= Binomial class, GD = Growth Defect class.

Lastly, we determined the number of EGs identified with bowtie (Gumbel or HMM) *versus* FiTnEss, which returned a FWER stringent list included in the FDR results (see Materials and methods section for a description of the statistical methods). As expected from previous results Gumbel identified very few EGs (205 EGs and 274 EGs) and around 50 % of gold standard EGs were present in these sets. HMM produced better results with over 90 % of gold standard EGs identified in the list obtained by considering both EGs and growth defect genes. FiTnEss, even with the stringent list of 358 EGs produced with FWER, contained more than 95 % of the GOLD_84 and 90 % of the GOLD_115 EGs. Reducing stringency with FDR yielded an EGs list of 609 genes containing 100 % of the gold standard EGs (Table 2). FiTnEss was the only method capable of identifying all the expected genes previously identified as essential and appeared to return a reliable list of EGs. Overall, these findings showed that bowtie_HMM and FiTnEss FDR were recommended for our Tn-Seq analysis.

**Table 2:** Number of EGs identified with Gumbel, HMM and FiTnEss methods.

|  | Number of EGs | GOLD_84<sup>a</sup> | GOLD_115<sup>a</sup> |
| --- | --- | --- | --- |
| **GUMBEL_EG** | 205 | 48,81 | 46,09 |
| **GUMBEL_E+EB** | 274 | 52,38 | 53,91 |
| **HMM_EG** | 184 | 38,10 | 37,39 |
| **HMM_Gs+GD** | 565 | 91,67 | 90,43 |
| **FiTnEss_FWER** | 358 | 95,24 | 89,57 |
| **FiTnEss_FDR** | 609 | 100,00 | 100,00 |

<sup>a</sup>The percentage of the reduced reliable list of gold standard EGs GOLD\_84 and GOLD 115.

### Identification of the EGs specific to PA14Δ*oprD*

As FiTnEss produced a list of EGs for one condition, we intersected the set of EGs provided by FiTnEss FDR from PA14 WT and PA14Δ*oprD* to obtain the specific EGs of the OprD-deficient PA14 strain in LB. In the genes obtained with bowtie_HMM, we identified genes exhibiting statistically significant variability of insertion counts across PA14 WT and PA14Δ*oprD* (conditionally EGs) either by three statistical methods (Anova, Resampling and Zinb) available in the TRANSIT suite to determine, or by intersecting as performed for FiTnEss (see Material and Methods and Supplementary data for more details).

As intersections might lead to false positive, we filtered out the genes specific to one condition (WT or Δ*oprD*) that are not classified as non-essential in all samples in the other condition (Δ*oprD* or WT, respectively). This filter reduced the number of genes for FiTnEss but did not impact the number of genes obtained with bowtie_HMM. The total of conditionally EGs for PA14Δ*oprD* and PA14 WT were 31 (bowtie_HMM_Anova), 49 (bowtie_HMM_Resampling), 218 (bowtie_HMM_Zinb), 197 (FiTnEss_FDR), 117 (FiTnEss FDR_FILTER) and 135 (bowtie_HMM_intersection). Only 3 genes were common between all the methods. The methods based on intersection (FiTnEss_FDR, FiTnEss FDR_FILTER and bowtie_HMM_intersection) shared more EGs among the dataset than those obtained with the statistical methods. By focusing on the shortest list of 117 EGs found by FiTnEss FDR_FILTER, we identified 30 conditionally EGs for PA14Δ*oprD* vs PA14 WT. As our 84 gold standard EGs (GOLD_84) were supposed to be essential for all PA strain in any culture medium, we confirmed they were not included in the aforementioned lists of conditionally EGs (only 3/84 genes were not absent in the set of genes obtained with bowtie_HMM). Moreover, we confirmed that the *oprD* gene was considered as EG in PA14Δ*oprD* but not in PA14 WT. Finally, we obtained a short but reliable strong set of 30 EGs in LB specific to PA14Δ*oprD* compared with PA14WT (Table 3).

**Table 3:** List of RPKM for the EGs determined to be specifics to PA14Δ*oprD*.

| Locus Tag | Name | RPKM PA14 WT | RPKM PA14Δ*oprD* |
| --- | --- | --- | --- |
| PA14_RS01270 | / | 23.666 | 2.484 |
| PA14_RS04505 | / | 20.633 | 0 |
| PA14_RS05905 | *iscX* | 35.395 | 0.6319 |
| PA14_RS09230 | *cpxR* | 7.8689 | 1.2182 |
| PA14_RS11895 | *tsi2* | 67.401 | 0.8142 |
| PA14_RS12140 | *bqsS* | 80.294 | 0.8071 |
| PA14_RS13385 | / | 198.69 | 30.934 |
| PA14_RS13560 | *pvdG* | 62.052 | 10.998 |
| PA14_RS13665 | / | 25.501 | 6.6364 |
| PA14_RS13690 | / | 23.72 | 1.7321 |
| PA14_RS14055 | / | 3.8567 | 0 |
| PA14_RS14990 | / | 26.746 | 3.0256 |
| PA14_RS15035 | / | 8.5298 | 0 |
| PA14_RS15170 | *opdO* | 101.93 | 12.968 |
| PA14_RS15565 | / | 6.0981 | 0 |
| PA14_RS15605 | *mexX* | 19.539 | 0.1067 |
| PA14_RS15660 | *fahA* | 21.369 | 3.132 |
| PA14_RS15800 | *exaE* | 23.264 | 3.0936 |
| PA14_RS17305 | *pscS* | 27.58 | 3.5705 |
| PA14_RS18035 | *ccoO* | 48.767 | 13.558 |
| PA14_RS18045 | *ccoP* | 30.46 | 8.4445 |
| PA14_RS19615 | / | 31.314 | 1.7252 |
| PA14_RS22095 | *pdxJ* | 93.984 | 6.8983 |
| PA14_RS22870 | / | 17.336 | 0 |
| PA14_RS25565 | *pcnB* | 33.939 | 16.314 |
| PA14_RS25580 | *panC* | 25.796 | 4.9059 |
| PA14_RS27060 | *aceE* | 63.29 | 17.359 |
| PA14_RS27920 | / | 21.391 | 0.8821 |
| PA14_RS29395 | *prfH* | 51.865 | 0.1033 |
| PA14_RS30300 | / | 137.36 | 0 |

Most of these genes were not annotated and therefore have as yet unknown functions. Since OprD is a specific porin that imports small basic amino acids (arginine, lysine, histidine and ornithine) into PA, we focused on genes related to the metabolism of those basic amino acids. Two genes leading to precursors of the tricarboxylic acid (TCA) cycle were identified: *panC* and *fahA. panC* encodes a pantoate--beta-alanine ligase (also known as pantothenate synthetase) that produces (R)-Panthothenate, which plays a role in the production of Coenzyme A (22). *fahA* encodes a fumarylacetoacetase that produces fumarate, which is one of the key components of the TCA cycle (23). As the TCA cycle provides precursors of certain amino acids (24,25), and gluconate is used in the Entner-Doudoroff pathway that produces pyruvate, needed for TCA cycle (26), we hypothesized that lacking *panC* or *fahA* precludes PA14 from compensating such a nutritive defect, in which the import of arginine, lysine, histidine and ornithine is diminished due to a non-functional OprD porin (Δ*oprD*).

### Silencing *panC* but not *fahA* affects PA14Δ*oprD* growth

We sought to confirming biologically the essentiality of these two genes for PA14Δ*oprD* grown in LB. To this end, we first attempted to carry out clean deletions of *panC* and *fahA*. We succeeded in constructing PA14 WT as well as PA14Δ*oprD* mutants lacking *fahA.* Neither of these two FahA-deficient isogenic strains showed any growth defect when grown in LB media. *fahA* was therefore not an essential gene for growth in LB for PA14 (WT and Δ*oprD*). In contrast to the *fahA* gene, despite numerous attempts, we were unable to cleanly delete *panC* in PA14 WT nor in PA14Δ*oprD.* The inability to delete *panC* in PA14 WT and PA14Δ*oprD* suggested its essential character for LB growth. However, we could not rule out a bias since we used the same media for deletion and Tn-Seq.

Consequently, we aimed at gradually silencing *panC* to determine whether its low or near-zero expression could prevent the growth of PA in LB. To this end, we employed the CRISPR Inhibition method using a Mobile-CRISPRi vector, which embedded a gene-specific sgRNA and a catalytically dead *Cas9* inducible with arabinose (27,28), to control gene expression. CRISPRi isogenic strains were constructed to silence *panC* in PA14 WT and PA14Δ*oprD*, but also *fahA*, to confirm our previous results. Both CRISPRi PA14 WT and PA14Δ*oprD* were compared with their isogenic control strains carrying a CRISPRi module with a non-specific sgRNA that did not bind to the PA chromosome.

Although the growth curves were quite similar for the isogenic strain grown in LB +/- 0.1% arabinose (no and weak *panC* silencing), we observed a growth defect for the OprD-mutant strain when *panC* was deeply silenced at the maximal non-toxic concentration of arabinose (1%) (Fig. 2A) (27). To a lesser extent, such a growth defect was also observed for PA14 WT. To complete our LB growth analysis when *panC* was silenced, we determined the Maximum Growth Rate (MGR) of each growth curve. As shown in Figure 2B, at 1% arabinose, we evidenced a decrease in MGR in *panC*-silenced PA14 WT and PA14Δ*oprD* (4.25.10^-4^s^-1^ and 3.64.10^-4^s^-1^, respectively) compared to their respective control (5.10.10^-4^s^-1^ and 5.09.10^-4^s^-1^, respectively). Interestingly, silencing *panC* had a more negative impact on bacterial growth of PA14Δ*oprD* than PA14 WT in LB.

**Figure 2.**
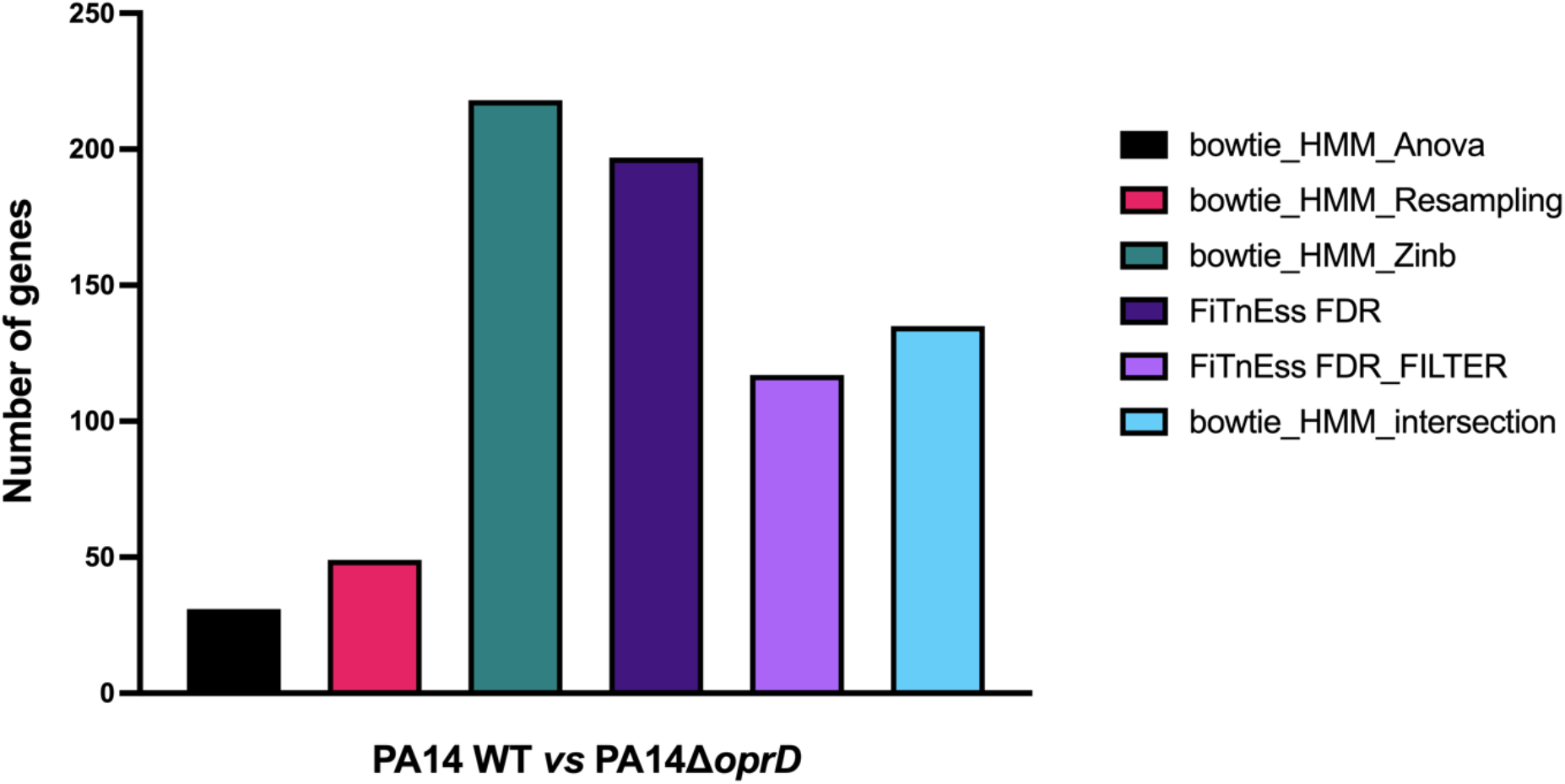
Number of genes conditionally essential in LB for PA14Δ*oprD* compared to PA14 WT. Comparison of the numbers of GEs only in one of the two PA14 strain tested obtained with different analysis methods.

**Figure 3:**
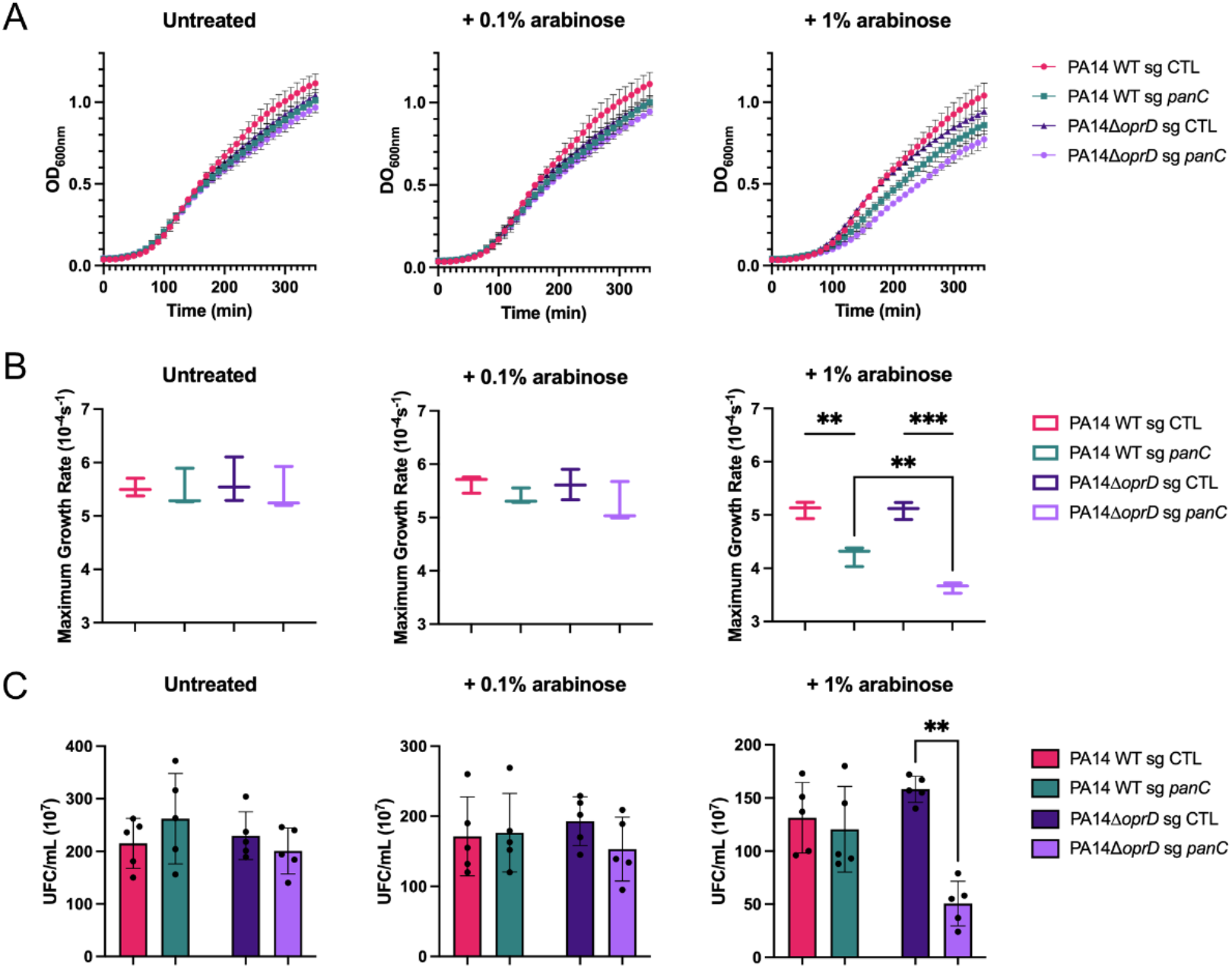
Biological verification of *panC* essentiality in PA14Δ*oprD* shows reduced growth and biomass. **(A)** Growth curves of PA14 WT sg CTL, PA14 WT sg panC, PA14Δ*oprD* sg CTL and PA14Δ*oprD* sg panC (pink, green, dark purple and light purple respectively) were realized in Tecan. LB media supplemented were supplemented with various concentration of arabinose (0.1% or 1%) for 6 hours (n=3). **(B)** Maximal Growth Rate of the PA14 CRISPRi strains for the Tecan growth curves. Maximum Growth Rate (MGR) was determined as the maximum value of the derivative of the logOD_600_ using R software. Statistics were achieved by unpaired t-test: ***, *p* < 0.001; **, *p* < 0.01. **(C)** Colony count of the PA14 CRISPRi stains after culture LB media supplemented with various concentration of arabinose (0.1% or 1%) for 6 hours. All data represent mean values of 5 independent biological replicates, and error bars indicate standard deviation. Statistics were achieved by Mann-Whitney test. Asterisks indicate values that are significantly different as follows: \**p* < 0.05; \*\**p* < 0.01; \*\**p* < 0.001; ns (Not significant).

Next, using colony-forming unit counting, we determined cell viability according to whether the *panC* gene was silenced or not. On one hand, we found no difference in CFU for PA14 WT (sg *panC*) compared with its control (sg CTL) at 0, 0.1% and 1% of arabinose (Fig. 2C). On the other hand, we found no difference in CFU for PA14 Δ*oprD* (sg *panC*) compared with its control (sg CTL) at 0 and 0.1% arabinose, but when *panC* was deeply silenced (1% of arabinose), we evidenced a decrease of 68% of CFU compared to its control (sg CTL).

Using these approaches to assess the growth of *fahA*-silenced PA14 WT and PA14 Δ*oprD*, we confirmed that *fahA* was not essential for growth in LB of the PA14Δ*oprD* strain. Indeed, we found no difference in terms of growth curves, MGR and CFU counting between the sg *fahA* strains and their sg CTL isogenic control strains (Fig. S2).

Overall, these results showed that silencing *panC* had a negative impact on the growth of the OprD-mutant PA14 strain in LB. This allowed us to define *panC* as a “growth defect” gene. However, given that the CRISPRi method is not equivalent to a clean deletion, it is likely that the persistence of a very slight expression of *panC* explained why we still harvested CFUs and therefore were unable to classify unambiguously this gene as an EG for PA14 Δ*oprD in vitro* (LB).

### Lack of uptake by OprD does not explain the need of *panC* for *in vitro* growth

We assessed whether compensating the decrease of basic amino acids produced by the TCA could overcome the growth defect in PA14 WT but not in PA14 Δ*oprD* when *panC* was silenced. Using minimal media (M9 minimal salt) supplemented with arginine, lysine, histidine and ornithine but also with gluconate (also imported by the OprD porin), we counted CFUs on agar plates after CRISPRi silencing of *panC* (1% arabinose, for 6h). Although we observed less viable PA14 cells for both *panC*-silenced PA14WT and PA14Δ*oprD* (Sg *panC*) than for their respective controls (Sg CTL) when supplemented with amino acids (Fig. 4), no statistical difference was found between PA14 WT (green bar) and PA14Δ*oprD* (purple bar). Such a supplementation was not sufficient for PA14 WT to cope with the lack of *panC* and provide better growth than the OprD mutant PA14, which cannot benefit from such supplementation. These results showed that lack of import by the OprD porin did not explain why silencing *panC* led to an *in vitro* growth defect of PA14 Δ*oprD*.

**Figure 4:**
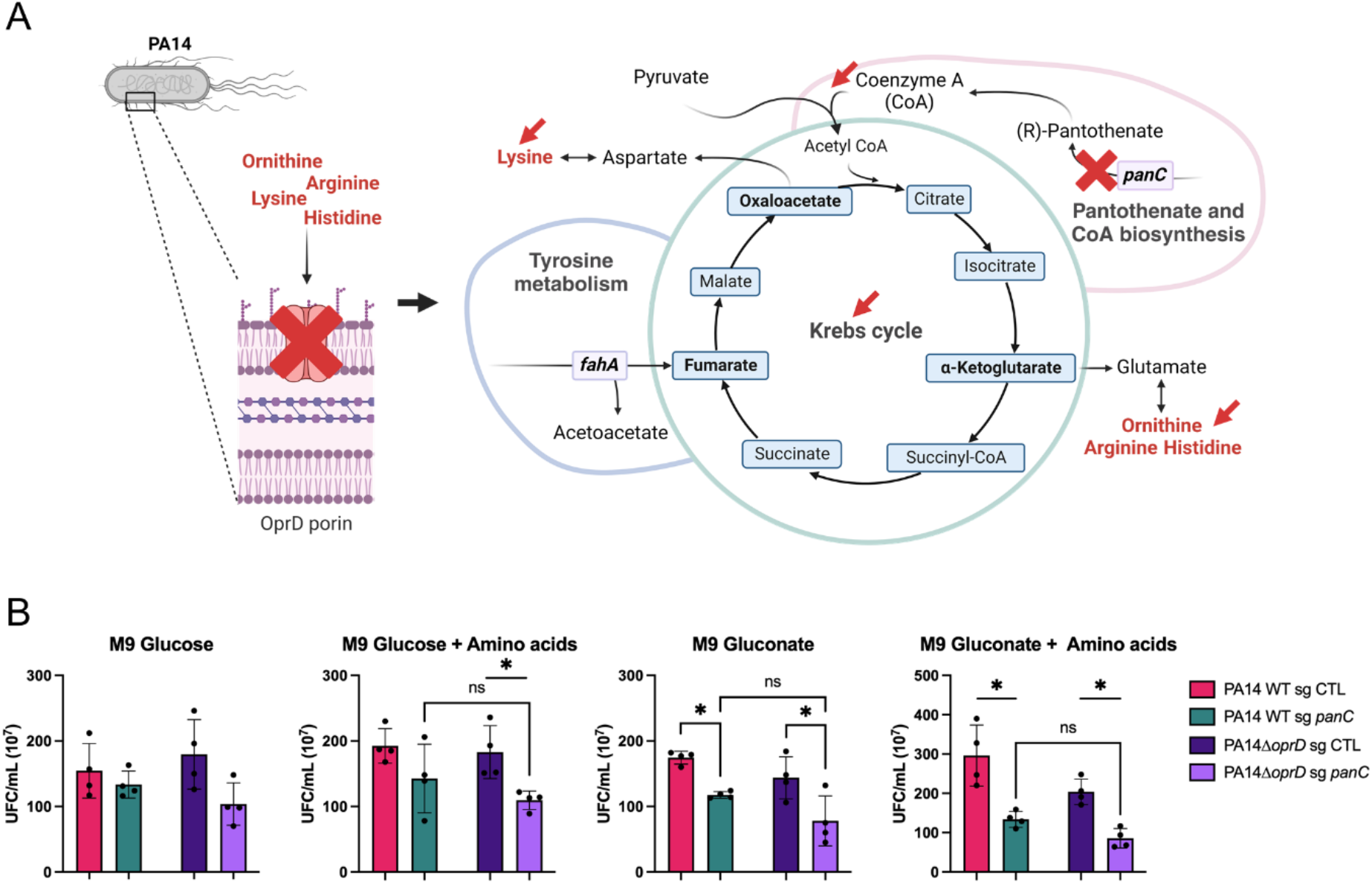
Lack of import linked to OprD porin loss did not cause *panC* essentiality for the PA14Δ*oprD* strain. **(A)** Hypothesis of the consequences for PA14Δ*oprD* of the inhibition of *panC*. Red arrows indicate the supposed diminutions observed when *panC* is not functional. Created with BioRender.com. **(B)** Colony count of the PA14 CRISPRi stains after culture with 1% arabinose in Minimal Media M9 supplemented with glucose or gluconate, with or without the 4 amino acids using the OprD porin (Arginine, Lysine, Histidine and Ornithine) for 6 hours. All data represent mean values of 4 independent biological replicates, and error bars indicate standard deviation. Statistics were achieved by Mann-Whitney test. Asterisks indicate values that are significantly different as follows: \**p* < 0.05; \*\**p* < 0.01; \*\**p* < 0.001; ns (Not significant).

### Silencing *panC* impacts the FAS pathway in OprD-deficient PA

Given that lacking the *panC*-encoded pantothenate synthetase could preclude CoA production, which is needed for fatty acids (FA) production (29), we hypothesized that silencing *panC*, could lead to deregulate FA synthesis (FAS) and affect the membrane structure.

FA play an important role in membrane structure, modulating membrane fluidity. The more unsaturated FA present, the greater the membrane fluidity. It has been reported that *sigX* regulates the expression of *cmpX* and various enzymes involved in FAS by balancing the ratio of saturated and unsaturated FA. Overexpression of *sigX* increased short FA, the *fabD*, *fabG*, *fabY* and *fabZ* genes are each involved in different stages of the FAS pathway, and then membrane fluidity (30–33) (Fig. 5A). Accordingly, we assessed the expression of these genes in the context of silencing *panC* in the isogenic PA14Δ*oprD* and PA14 WT strains. As shown in Figure 5B, we found no difference in *sigX* and *cmpX* expression between the two CRISPRi control strains (1.03 and 1.03-fold change respectively for PA14Δ*oprD* sg CTL compared to PA14 WT sg CTL). Therefore, loss of the OprD porin did not appear to impact *sigX* and *cmpX* expression. However, we highlighted that *sigX* expression was drastically decreased upon *panC* silencing in PA14 WT sg *panC* and PA14Δ*oprD* sg *panC* compared to PA14 WT sg CTL (0.47 and 0.26-fold change respectively), with slightly lower expression in the OprD mutant strain (Fig. 5A). We noted a trend towards the decreased expression of *cmpX* in PA14Δ*oprD* sg *panC* with a non-significative 0.54-fold-change. Our results were found to be agreement with a less fluid membrane.

**Figure 5:**
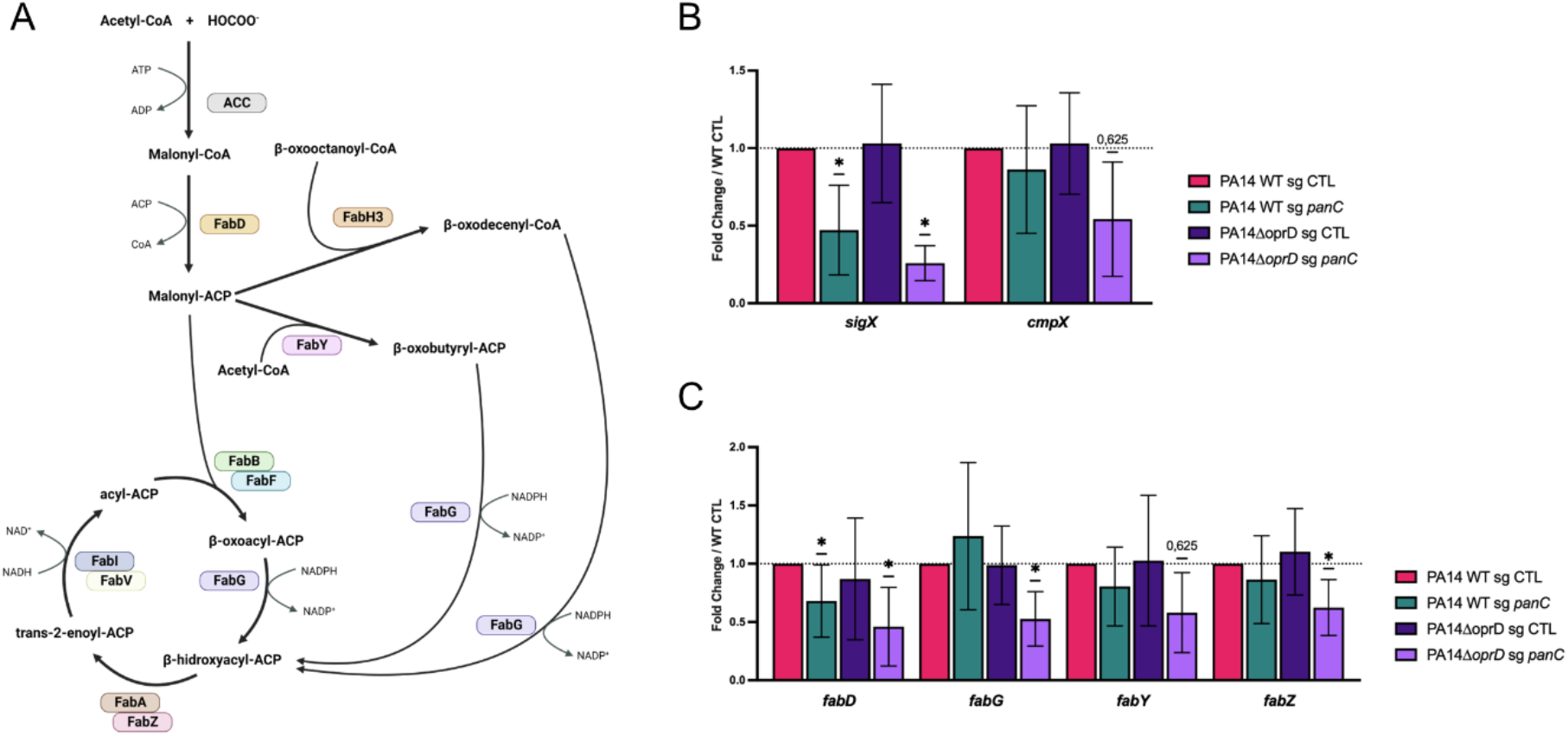
Silencing *panC* deregulates the expression of FAS genes. **(A)** FA biosynthesis pathway. **(B)** Relative expression of *sigX* and *cmpX* in PA14 WT sg *panC* (dark green) and PA14Δ*oprD* sg *panC* (light purple) in comparison to their isogenic PA14WT and PA14Δ*oprD* strain carrying sg CTL (pink and dark purple respectively), exposed to 1% arabinose for 6 hours, normalized with *gyrB*. **(C)** Relative expression of *fabD*, *fabG*, *fabY* and *fabZ* in PA14 WT sg *panC* (dark green) and PA14Δ*oprD* sg *panC* (light purple) in comparison to their isogenic PA14WT and PA14Δ*oprD* strain carrying sg CTL (pink and dark purple respectively), exposed to 1% arabinose for 6 hours, normalized with *gyrB*. All data represent mean values of 6 independent biological replicates, and error bars indicate standard deviation. Statistics were achieved by Wilcoxon test. Asterisks indicate values that are significantly different as follows: \**p* < 0.05; \*\**p* < 0.01; \*\**p* < 0.001; ns (Not significant).

We did not find any significant change in *fabD, fabG, fabY* and *fabZ* expression in PA14Δ*oprD* sg CTL compared with PA14 WT sg CTL (0.87, 0.98, 1.02 and 1.10-fold change, respectively) (Fig. 5C). Although we showed a non-significant 0.58-fold change expression of *fabY* for PA14Δ*oprD* sg *panC* compared PA14 WT sg CTL, we found decreased expressions of *fabD, fabG* and *fabZ* (0.46, 0.52 and 0.62-fold change, respectively). For *fabD*, we also observed a decreased expression for PA14 WT sg *panC versus* its control (Fig. 5C). Overall, these results indicated a decrease in unsaturated FA synthesis when *panC* was silenced, which and could therefore affect the membrane homeostasis. This could explain the essential nature of the *panC* gene in the OprD-defective PA14 strain.

### Silencing *panC* reduces virulence of OprD-deficient PA against 16HBE human bronchial epithelial cells

Carbapenem-resistant OprD-deficient PA strains are a global threat as physicians may face therapeutical impasses. In light of our findings concerning the functional and metabolic alterations of PA14 Δ*oprD* upon silencing *panC*, we investigated whether a therapeutical strategy aiming at inhibiting *panC* could be an anti-virulence strategy for treating PA infections. We infected 16HBE cells with PA14 WT sg *panC* and PA14Δ*oprD* sg *panC*, as well as their isogenic CRISPRi control strains (PA14 WT sg CTL and PA14Δ*oprD* sg CTL) and monitored the 16HBE cell death for 48 hours by time-lapse microscopy. When setting up our 16HBE infection model, we found that 1% arabinose, for CRISPRi induction, precluded propidium iodide-based fluorescence monitoring as well as image analyses (Fig. S3). We assumed that 1% arabinose may reach a toxic concentration for bacteria delaying the time of cell death of 16HBE by all CRISPRi strains. Then, we induced CRISPRi upon exposure to 0.5% arabinose during 16HBE infection. To rigorously analyze the images obtained from time-lapse microscopy, we used 3 different techniques to compare three readouts for assessing 16HBE cell death by PA: (i) the 50% intensity time (IT50), (ii) the threshold time according to intensity (TTI) and (iii) the 50% of dead cells time (DCT50) (Fig. 6). We evidenced increased virulence for PA14Δ*oprD* sg CTL against 16HBE cells compared to PA14 WT sg CTL with lower TTI (11.74 *vs* 14.94h, respectively) and DCT50 (11.44 *vs* 17.94h, respectively) (Fig. 6C). These results were in agreement with previous studies (11–13). We found an increased IT50 when inhibiting *panC* in PA14 WT (27.4h for sg *panC vs* 21.9h for sg CTL) while TTI and DCT50 readouts were not statistically increased. This could be explained by the late fluorescence intensity associated with bacterial death that created a haze interfering with the intensity measurements, which was therefore erroneously considered as delayed. IT50 turned out to be the least reliable readout, which reinforced our strategy of analyzing cell death in terms of IT50, TTI and DCT50. Such a discrepancy led us to draw no conclusion on the effect of *panC* inhibition in PA14 WT. In the case of PA14Δ*oprD*, based on the three readouts, we found that the time of 16HBE cell death following infection was greatly extended when *panC* (PA14Δ*oprD* sg *panC*) was silenced compared with its isogenic control (PA14Δ*oprD* sg CTL) (26.5h *vs* 11.4h for DCT50; 16.9h *vs* 11.7h for TTI; and 26.8h *vs* 19.7h for IT50). Moreover, silencing *panC* in PA14Δ*oprD* also showed extended cell death time compared to PA14 WT sg CTL (respectively 16.9h vs 14.9h with TTI and 26.5h vs 17.9h with DCT50). Overall, we showed that *panC* inhibition reduced the virulence of the carbapenem-resistant OprD mutant PA14Δ*oprD*.

**Figure 6:**
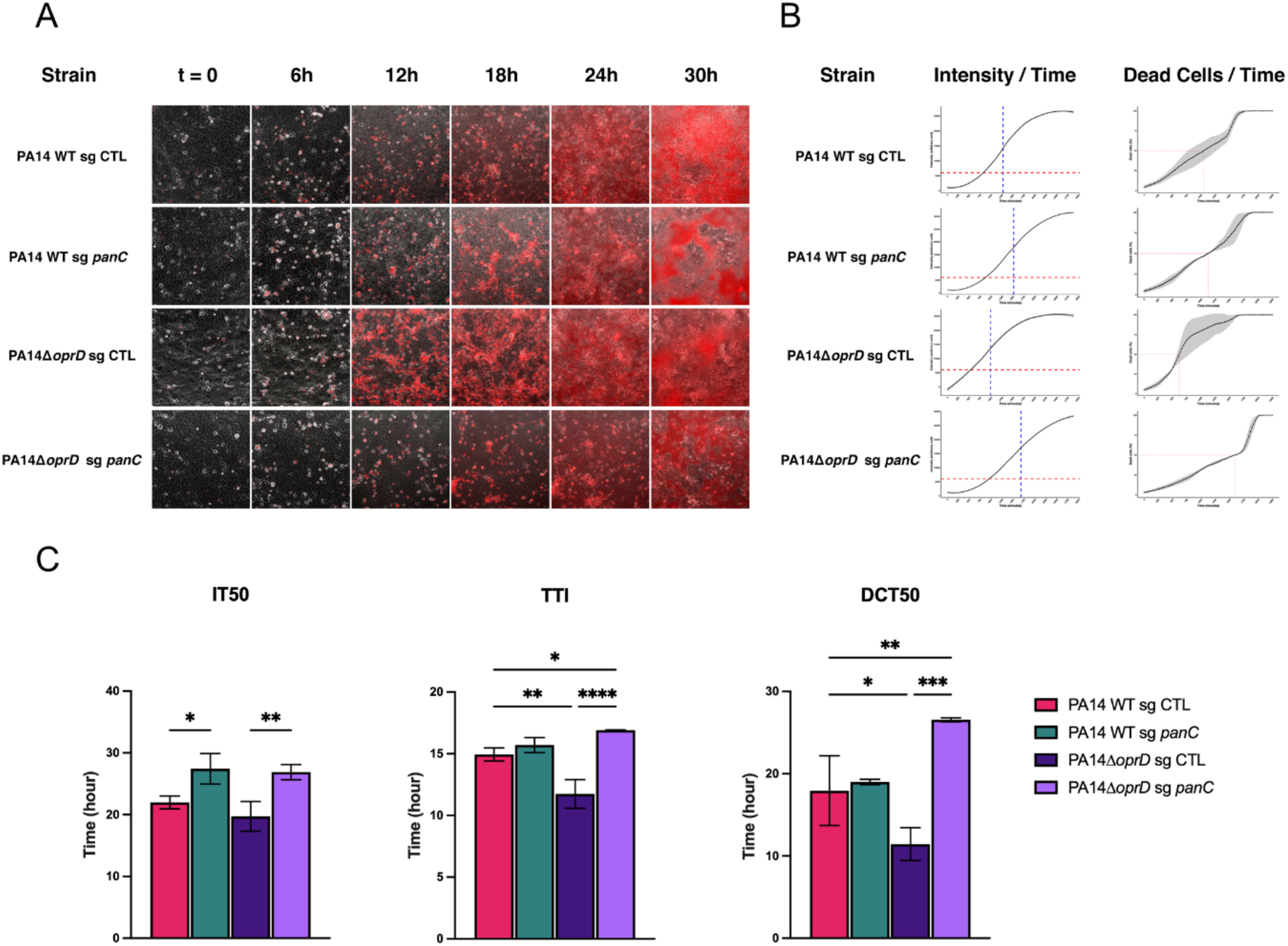
Silencing *panC* reduced virulence of OprD-deficient PA against 16HBE cells. **(A)** Kinetics of 16HBE cell death following infection with CRISPRi isogenic OprD-producing (WT sg CTL and sg *panC*) and its OprD-defective derivative PA14 strains (Δ*oprD* sg CTL and sg *panC*) for *panC* inhibition. Dead cells are marked with propidium iodide and appear red. **(B)** Curve of intensity over time corresponding to the well shown in (A). The red dotted line corresponds to the threshold time according to intensity (TTI) that is the time at which the intensity measured crosses the arbitrary threshold set at 6000 (arbitrary units). The blue dotted line corresponds to the 50% intensity time (IT50). Percentage of dead cells over time is represented on the right. Red lines indicate the time corresponding to 50% dead cells time (DCT50) for each strain. Standard deviations are displayed in grey. **(C)** Average time for cell death for PA14 WT sg CTL, PA14 WT sg *panC*, PA14Δ*oprD* sg CTL and PA14Δ*oprD* sg *panC* depending of the used readout (IT50 on the left, TTI in the middle and DCT50 on the right). All data represent mean values of 3 independent biological replicates, and error bars indicate standard deviation. Statistics were achieved by ordinary one-way ANOVA followed by Tukey’s multiple-comparison test. Asterisks indicate values that are significantly different as follows: \**p* < 0.05; \*\**p* < 0.01; \*\**p* < 0.001; ns (Not significant).

## Discussion

The loss of the OprD porin is the major determinant of resistance against the last resort carbapenem antibiotics (34). OprD loss has been described as a basis for enhanced fitness and virulence *in vivo* (11–13). Therefore, tackling carbapenem-resistant OprD-deficient PA is in line with the World Health Organization (5,35), which considers carbapenem-resistant PA as a critical priority for which new antimicrobial drugs or alternate therapies are needed. Following the example of many studies that have determined EGs PA14 WT in various condition (11,15,19,20), we used Tn-Seq to determine, for the first time, the specific EGs of PA14Δ*oprD* in LB. After evaluating the best bioinformatic method to analyze our Tn-Seq data, we found an EG for *in vitro* growth of PA14Δ*oprD* that was likely to be a potential target for alternate therapies.

A large number of algorithms have been developed to analyze Tn-Seq data, including HMM-based methods for identifying essential sites and regression analyses using gene saturation or series of consecutive empty sites. These approaches were implemented in tools such as TnseqDiff (36) ESSENTIAL (37), Magenta (38), Tn-seq Explorer (39), ARTIST (40), TRANSIT (18), TSAS (41). FiTnEss, was introduced to define the core essential genome of PA, as another statistical method developed to identify EGs (15). TRANSIT is the most popular tool and includes a high number of tools for analyzing Tn-Seq data, from pre-processing to gene enrichment. One of the main limitations of available studies is the absence of a gold standard set of EGs to evaluate the analysis process. Indeed, many studies determined EGs for PA based on their own reference and not on a set of reference genes that could be used to initially validate any method to be considered. Thus, (i) by establishing two sets of gold standard EGs (GOLD_115 and GOLD_84), we concluded that Bowtie_HMM and FiTnEss enabled us to identify reliable EGs of each PA14 WT and PA14Δ*oprD*, and (ii) by intersecting the EGs (FiTnEss FILTER and Bowtie_HMM_intersection), we identified a strong, short but reliable set of 30 EGs in LB specific to PA14Δ*oprD* versus PA14WT. We anticipate that making our GOLD_115 and GOLD_84 publicly available will help the scientific community for future Tn-Seq analyses.

Among the EGs for *in vitro* growth in LB of PA14Δ*oprD*, we focused on two genes, *fahA* and *panC*, linked to the TCA pathway. It is worth noting that despite all the bioinformatics precautions described above, the biological validation of an EG remains an imperative. We invalidated *fahA* since its PA14Δ*oprD* clean deleted mutant, among other things (see results) was viable *in vitro*. For *panC*, we failed to delete it in PA14Δ*oprD* but we showed, using a CRISPRi approach, that when silenced, it impacted PA14Δ*oprD* with reduced growth in LB. Such a strategy did not fully inhibit gene expression, unlike clean deletion. Thus, although we could not affirm that *panC* is a *stricto sensu* EG, it could at least be considered a growth defect gene for PA14Δ*oprD* in LB.

*panC* codes for pantothenate synthetase that produces pantothenate, which is required for CoA production. Moreover, within cultures of *Escherichia coli* or *Salmonella* Typhimurium, it has been reported that pantothenate supplementation increased the synthesis of molecules requiring CoA intermediates (42–44). Hence, as CoA is needed in the TCA cycle, deregulation of CoA production by *panC*, in the pantothenate pathway, could impact the downstream effects of the TCA cycle (45,46). Our first hypothesis to link the EG *panC* to OprD was that the deregulation of the TCA could decrease arginine, lysine, histidine and ornithine synthesis, and thus could not compensate for the lack of import of these basic amino acids due to the loss of OprD. Using culture supplemented with these basic amino acids, we showed that the lack of uptake by OprD did not explain the essentiality of *panC* for *in vitro* growth. Given that, OprD is the second major porin present within the membrane of PA, OprF being the first one (10), and OprF has been previously described as playing a key role in maintaining PA membrane integrity (10), we questioned whether FAS impairment following TCA deregulation could have a highly deleterious impact on the membrane of the OprD-deficient PA14 strain. We found a decreased expression of the mechanosensitive ion channel-encoding *cmpX* gene and its regulator, the extracytoplasmic sigma factor *sigX* (47) in PA14Δ*oprD* when *panC* was silenced, which pointed to altered membrane structure. In line with these results, we showed a decrease in the expression of genes involved in FAS, particularly *fabD*, which is involved in an early stage of the FAS (32), and *fabG, fabY* and *fabZ*, which participate in FA elongation leading to unsaturated FAs (48). Taken together, these results indicated that the essentiality of *panC* in the OprD-defective PA14 strain was likely due to an alteration in membrane homeostasis consecutive to a decrease in unsaturated FA synthesis in the absence of *panC*.

While all organisms synthesize CoA, humans cannot synthesize pantothenate *de novo*, offering therefore a bacteria-specific target. Such a strategy was previously considered (49,50), but sidelined because antimicrobial susceptibility testing failed to show *in vitro* growth inhibition of bacteria treated with competitive inhibitors of pantothenate synthetase (51,52). It was suggested that either pantothenate synthetase was not susceptible to inhibition or its inhibition was irrelevant to bacterial survival and growth. However, it has been recently described that CoA synthesis was essential in *Toxoplasma gondii* tachyzoites because they were unable to salvage CoA or intermediates from the pathway (53). Thus, pantothenate synthesis has been proposed as a promising target for *T. gondii* persistence. On one hand, based on the essentiality of *panC in vitro* for PA14Δ*oprD*, we took into question the irrelevance of inhibiting the pantothenate synthetase. On the other hand, taking our results into account, these recent findings boosted our interest in targeting *panC* during PA infections as an alternate therapeutical strategy in the context of antimicrobial resistance. PA is a major pathogenic bacterium involved in often fatal respiratory tract infections in cystic fibrosis and chronic obstructive pulmonary disease patients but also critically ill patients hospitalized in intensive care unit. PA penetrates through the airway tract, and the human airway epithelium is the physical and functional barrier that prevents infection from inhaled pathogens (54). For this reason, we assessed the anti-virulence potential of *panC* inhibition in a 16BHE model of PA infection. We showed that silencing *panC* by CRISPRi extended the time of cell death by the carbapenem-resistant OprD-defective PA14 strain compared to that of the non-silenced strain.

To conclude, we introduced a gold standard dataset of EGs available to compare the metrics of different methods for Tn-Seq analysis. It allowed us to provide, for the first time, the EGs *in vitro* of PA14Δ*oprD.* The reason of essentiality of one of them, *panC*, was deciphered and we shed a new light on its inhibition as a target for treating carbapenem-resistant OprD-defective PA pulmonary infections. We are aware that our results represent only a first step, but we anticipate to further complete our findings by identifying the EGs of PA14Δ*oprD* during infection *in vivo* and searching for potential *panC* inhibitors.

## Material and methods

### Bacterial strains and growth conditions

The laboratory strain *P. aeruginosa* PA14 UCBPP (55) (and its derivatives were used in this study. The PA14 parent wild-type (WT), mutant lacking OprD (Δ*oprD* by clean deletion) and the OprD mutant complemented with *oprD* under the control of its native promoter strains were gifts from D. Skurnik (11,12). CRISPRi strains were made using two plasmids pJMP1039 and pJMP1237, that were a gift from Carol Gross & Jason Peters & Oren Rosenberg (Addgene plasmid # 119239; http://n2t.net/addgene:119239; RRID:Addgene_119239 - Addgene plasmid # 119262; http://n2t.net/addgene:119262; RRID:Addgene_119262) with *E.coli MFD* strain as donor. CRISPRi strains were made for silencing two gene, *panC* and *fahA* in PA14 WT and PA14Δ*oprD*, and a control strain was made using a non-targeting spacer against PA14 DNA. List of the strains used for this work are referenced in table S1 and plasmids in table S5. Bacterial cultures were incubated at 37 °C on Lysogeny Broth (LB) (BD Difco™) or M9 Minimal Salts, 5x (BD Difco™) when needed under agitation at 250 rpm.

### PA14 Δ*oprD* TnBank preparation

To construct the TnBank, PA14 Δ*oprD* was grown overnight in lysogeny broth (LB) and *E. coli* SM10λpir/pSAM_DGH was grown overnight in LB + gentamicin 15μg/mL at 37°C at 250 rpm. Equal volumes of both cultures were mixed and centrifuged for 5 min at 3000 rpm. Pellets were resuspended with 200 µL of water for two washes, and finally resuspended in 200 µL of LB. Mattings for conjugation were performed by plating mixed bacteria in spots onto LB agar for 3 hours at 37°C to transfer pSAM_DGH into PA14 Δ*oprD*. The transposon embedded in the plasmid was randomly inserted into a single TA site per PA14 Δ*oprD* bacterium, inactivating the gene wherever it was found. The spots were then resuspended in LB and mutants selected on LB + gentamicin 30 μg/mL + irgasan 25 μg/mL at 37°C for 24 to 48 hours. All colonies (around 300,000 mutants) were scrapped and conserved in cryotubes, which were finally pooled in 60 mL of LB with gentamicin 20 μg/mL and cultured for 2 hours at 37°C at 250rpm. The resulting PA14 Δ*oprD* Tn Bank was dispensed into 1mL cryotubes containing glycerol for storage at −80°C.

### Tn-Seq samples preparation

TnBanks samples were prepared as described previously (11–13). Briefly, TnBanks were grown overnight in 5mL of LB at 37°C at 250 rpm. Cultures were diluted to OD_600_=1 and centrifuged 15 min at 3000 rpm. Bacterial DNA was extracted with QIAamp^®^ minikit (Qiagen) and digested with MmeI enzyme for 4 hours at 37°C. Samples were then purified with QIAquick^®^ PCR Purification kit (Qiagen) and run on 2% agarose gel to only retrieve fragments between 1500 and 2000 bp, which were then purified with QIAquick^®^ Gel Extraction kit (Qiagen). DNA was next ligated to Illumina adapters and amplified by PCR to obtain libraries, which were then sequenced using a NextSeq500.

### Pre-processing steps

Reads were collected in a single file per sequencing run. To filter out PhiX reads, the reads of an Illumina technology control were mapped against the PhiX sequence with bowtie2 (56) and the resulting unmapped reads were retained. PhiX reads accounted for 24% and 23% respectively in each run. The reads contained the P2 adapter, then the sample-specific barcode (6 bp), 16-18 bp of the PA genome followed by the reverse complement of the 30-nucleotide P1M6GAMmeI sequence. (ACAGGTTGGATGATAAGTCCCCGGTCTATC). To identify the reads from each sample and extract the 16-18 bp of PA for downstream analysis, cutadapt (57) was used with this command line: cutadapt -g “sample_BARCODE(6nt);min_overlap=6;e=0…ACAGGTTGGATGATAAGTCCCCGGTC TATC;min_overlap=30;e=0” --discard-untrimmed -m 16 -M 18 -j 0 -o output.fastq noPhiX.fastq.

This command line extracted the subsequence with a size between 16 (-m 16) and 18 nucleotides (-M 18) that lay exactly between the sample barcode and the P1M6GAMmeI sequences in error-free reverse complement (e=0). At the end of this step, a Fastq file was obtained for each sample containing only the 16-18 bp sequences from PA genome.

### Read mapping and reads-counts file

Two mappers were used in this study using the trimmed fastq files: BWA (16) and bowtie (17). BWA version 0.7.12 was directly used with the TRANSIT v3.2.3 tpp command on the PA UCBPP-PA14 genome sequence (NC_008463.1), producing a .wig file. Bowtie 1.1.2 was used with the option -v 0 (no mismatches) -m 1 (report only unique mapping) producing a sam file. Scripts from https://github.com/SuzanneWalkerLab/5SATnSeq (sam_to_tabular.py and make_wig.py) were used to obtain a .wig file from the alignment file. FiTnEss took a sam file as input and returned a tally file with the script “tally.py”. Then the tally file is used with FitNess function in R with defaults parameters but manually updating the reference genome to NC_008463.1.

### Statistical methods to detect essential genes

The TRANSIT suite offers several statistical methods to detect essential genes. HMM proposes a 4-state model allowing for detecting regions as essential (E), non-essential (NE), growth-defect (GD) and growth-advantage (GA). The Gumbel method makes a call based on the longest consecutive sequence of TA sites without insertion into genes and calculates the probability of this using a Bayesian model. The important features of FiTnEss are that (i) it evaluates genes (rather than individual TA sites or stretches of TA sites) and (ii) it uses a simple two-parameter model to capture all the salient features of the data (the simple model having greater statistical power). To vary the stringency, two different levels of multiple testing adjustment can be applied: one with maximal stringency, which provides the most reliable set of essential genes [family-wise error rate (FWER)], to identify genes with few or no sequencing reads, and the other with high stringency but slightly relaxed [false discovery rate (FDR)] to identify genes that are statistically significant but contain a low number of reads. Genes with an adjusted *P-*value <0.05 in the replicates were predicted to be essential.

### Methods for EGs determination (software and normalization)

Firstly, the impact of software parameters was studied for all the methods. The -r parameter is available in the TRANSIT suite and allows replicates to be processed by averaging or summing the reads counts in the datasets. By default, read counts are summed for Gumble and averaged for HMM. This should not have effect on Gumbel, apart from potentially affecting spurious reads. For HMM with regular datasets (i.e. mean-read count > 100) it is recommended to average the read counts. For sparse datasets, it is recommended to sum the read counts, which can produce more accurate results. The HMM implementation suggested to use the loess option, which eliminates any bias linked to genomic position. Next, we investigated the impact of normalization. Totreads normalizes datasets over the total read counts and scales them to have the same mean over all counts; TTR (Trimmed Total Reads), normalizes over the total read counts (like Totreads), but trims top and bottom 5% of read counts. This is the recommended normalization method for most cases as it has the benefit of normalizing for difference in saturation in the context of resampling. Nzmean normalizes datasets to have the same mean over the non-zero sites. Quantile normalizes datasets using the quantile normalization method described by Bolstad et al. (58). Betageom normalizes the datasets to fit an “ideal” Geometric distribution with a variable probability parameter *p*. Especially useful for datasets that contain a large skew. Zinfnb fits a zero-inflated negative binomial model, and then divides read-counts by the mean. Finally, with nonorm parameter no normalization is performed.

### Methods for conditional essentiality identification

Resampling, proposed in TRANSIT, is based on a permutation test, and determines the read-counts that are significantly different from one condition to another. The ANOVA method, also available in TRANSIT, performs a one-way ANOVA test for each gene under all conditions. It takes into account the variability of normalized counts between TA sites and between replicates, to determine whether differences between mean counts for each condition are significant. Finally, the TRANSIT suite includes the Zero-Inflated Negative Binomial (ZINB) method, which can be applied to two or more conditions at once. The TRANSIT authors showed that the ZINB method typically identifies 10-20% more varying genes than resampling (and significantly outperforms ANOVA in detecting significant variability between conditions).

From results obtained from EGs identification methods, e.g. FiTnEss FDR or HMM_Es+GD, it was possible to intersect the lists of two conditions to obtain the EGs specific to each condition. Inspecting the results, we decided to add a filter to these sets of genes filtering out the essential genes in one condition which are not classified as non-essential for all samples of the other condition. Indeed, this case can potentially represent false positives, as the gene can be discarded from the list of essential genes simply because it is missing in one sample.

### EGs annotation

All the genes were annotated and clustered in function class using available resources from different databases like BioCyc Genome Database Collection (https://biocyc.org/PAER208963/organism-summary), The *Pseudomonas* Genome Database (https://pseudomonas.com/) (59) or GenomeNet (https://www.genome.jp/dbget-bin/www_bget?gn:pau).

### Deletion mutant construction

Deletions of genes in PA14 Δ*oprD* strain were performed using the replacement vector pEXG2 (60). Upstream and downstream fragments of ∼ 500pb flanking genes were amplified by PCR from PA14 genomic DNA using overlapping primers. Linearized fragment of pEXG2 was also obtained by PCR. The 3 fragments were ligated using the Gibson Assembly® Cloning Kit (New England Biolabs). The created pEXG2::Δ*fahA* and pEXG2::Δ*panC* were transformed into *E. coli* DH5α and positive clones selected on LB agar plates containing 15 mg/L of gentamicin. To transfer the plasmid in PA14 strain, triparental conjugation were performed between donor strain *E. coli*/ pEXG2::Δ*fahA* or pEXG2::Δ*panC*, recipient strain PA14 WT and PA14Δ*oprD* and helper strain *E. coli* HB101/pRK2013. The PA14 mutants were selected using LB agar mediums supplemented with irgasan 25mg/L and gentamicin 75mg/L. The merodiploid gentamicin-resistant PA14 strain was then cultured in LB broth containing 75 mg/L of gentamicin to exponential growth and streaked onto LB agar plates containing 12% sucrose for the allelic exchange. Sucrose-resistant colonies were tested to confirm gentamicin susceptibility, indicating excision from the genome of the pEXG2 backbone by double-cross-over event and thus gene replacement. Gene deletions were verified by PCR and sequencing.

### Construction of Mobile-CRISPRi vector and strains

The oligonucleotides used to create the sgRNA were designed as described by R.W. Ward (https://github.com/ryandward/Pseudomonas_sgRNA). Construction of vectors and strains was performed following published protocols (28) with slight adjustments. Briefly, annealed oligonucleotides were ligated to the BsaI-digested pJMP1237 plasmid. Such vectors and pJMP1039 were transferred in *E.coli* MFD DAP*-* strains by electroporation. Mobile-CRISPRi strains were constructed by tri-parental mating with PA14 WT or PA14Δ*oprD*. *E. coli* donor strains for the plasmids (*E.coli* MFD DAP-::pJMP1039 and *E.coli* MFD DAP-::pJMP1237 with chosen spacer) were grown overnight in LB + 300µM DAP + 100µg/mL ampicillin. The PA14 recipient strain were grown overnight in LB. Two milliliters of each culture were centrifuged at 7000 *x g* for 2min. Pellets were resuspended together in 1mL of LB and centrifuged and washed in LB two more times. Final pellets were resuspended in 200µL of LB and streaked on a cellulose filter and incubated overnight at 37°C. Filters were resuspended in 5mL of LB and vortexed. Pellets were resuspended in 200µL of LB and plated for selection on LB + gentamicin 75μg/mL. Vector integration were then verified by PCR and sequencing.

### Gene silencing with CRISPRi

Strains transfected with the constructed vectors were cultured overnight at 37°C in LB at 250 rpm. Cultures were adjusted to OD_600_=0.05 and cultured if needed with 0.1% or 1% arabinose either in 96 transparent flat bottom well plate using a SPARK^®^ multi-mode microplate reader (Tecan^®^) with 200µL per well for 24 hours at 37°C and 250 rpm and DO_600_ lecture every 10 minutes, or in a classic culture tube for 6 hours at 37°C and 250 rpm that afterward were diluted at 10^-6^ and spread on LB agar to determine survival by colony count after 24h at 37°C. For growth curves realized in SPARK^®^ multi-mode microplate reader, Maximum Growth Rate were determined following R script published in G. Royer paper (61). Results were analyzed by unpaired t-test.

### RT-qPCR

First-strand cDNA synthesis and quantitative real-time PCR was performed using the KAPA SYBR® FAST kit (CliniSciences) on the LightCycler® 480 (Roche Diagnostics) using the primers indicated in Supplementary Table S6. Gene transcript levels were normalized to *gyrB* as a housekeeping control gene. Gene expression levels were assessed using the 2^-ΔΔCt^ method (62,63) in respect to the MIQE guidelines (63). The relative fold-difference was expressed against the appropriate reference strains (PA14 WT CTL or PA14Δ*oprD*sg CTL). All experiments were performed as six independent replicates with all samples tested in triplicate.

### 16HBE cells infection

16HBE cell line was obtained from the American Type Culture Collection (Rockville, MD, USA). Cells were grown in a fully humidified atmosphere with 5% CO_2_ at 37°C. 16HBE cells were seeded on a Falcon 24-well plate (Corning, New York, United States), coated with type I collagen at 18.6 µg/mL (Sigma-Aldrich, Saint - Louis, United States), in Dulbecco’s Modified Eagle Medium (DMEM) GlutaMAX^TM^ (Gibco, Thermo Fisher Scientific, Waltham, United States) supplemented with 10% fetal bovine serum (FBS) and 1% penicillin (100 U/ml)/streptomycin (100 g/L). In each well, 350,000 cells were seeded on day 1 (D1), in DMEM GlutaMAX^TM^ medium with 10% FBS and 1% penicillin/streptomycin. At D3, when the cells were confluent, they were washed with Phosphate Buffered Saline (PBS) to remove antibiotics prior to infection with PA14 strains. For infection, 900 µL of FBS-free, antibiotic-free DMEM GlutaMAX^TM^ medium was added to 16HBE cells with propidium iodide and arabinose if required. Cells were infected with 100μL of PA14 culture adjusted to 10^6^ CFU/mL.

### Time-lapse microscopy

Infected 16HBE culture were placed in the incubation chamber of an inverted microscope (Axio Observer Z1, Zeiss^®^), at 37°C under 5% CO_2_. Using MetaMorph^®^ software (Molecular Devices), time-lapse microscopy was performed with a CMOS ORCA Flash 4.0 V2 camera (Hamamatsu, Japan), to record infection evolution by image acquisitions of the infected cells every 30 min for 72h (3 images/transwell). For cell death visualization, at each time point, a phase-contrast image and a red fluorescent image were recorded in parallel. Red fluorescence signal corresponding to propidium iodide necrotic dye was captured with a 545/25-nm excitation filter, a 570-nm beamsplitter, and a 607/70 bandpass emission filter. Composites images were processed using Image J software (National Institutes of Health, USA) and were then analyzed using three methods: 50% intensity time (IT50), time threshold according to intensity (TTI) and 50% dead cells time (DCT50).

### Fluorescence intensity analysis

IT50 is defined as the time required for the fluorescent signal to reach 50% of the maximal intensity, which correspond to the maximum intensity plateau obtained during infection. It was determined with log-logistics models by using “drc” package (https://CRAN.R-project.org/package=drc) on R Software (R 4.2.1 GUI 1.79, R Foundation for Statistical Computing, 2021). TTI is defined as the time required for the fluorescent signal to cross the threshold (ie exceeds background level). It was determined based on an intensity threshold calculated according to the spontaneous intensity of 16HBE cells over time, without bacteria, with and without the presence of 0.5% arabinose. Based on preliminary data, a threshold of 6000 (arbitrary units) was determined which correspond to an intensity point that cells without PA14 infection did not reach. Time corresponding to the threshold value was calculated using R Software.

### Cell Death Analysis

The Trainable Weka Segmentation (TWS) plugin, a machine-learning tool, was utilized for pixel-based classification (64). This plugin combines various image features with a wide array of machine learning algorithms from the Waikato Environment for Knowledge Analysis (WEKA) to categorize pixels into different class memberships. The procedural approach was as follows. No pre-processing was necessary. The TWS classifier was created using a dataset of 14 randomly selected images as the training data from a larger set of images. Region of Interest (ROI) selections were performed to classify IP positive cells as class 1 and background as class 2. Classifier was then trained and assessed for misclassified area. This training process was repeated multiple times to fine-tune the classifier. The training features used included Gaussian blur, the Hessian, the difference of Gaussian, and the Sobel filters. The classifier was subsequently validated using a dataset of 24 representative image sequences. For each of the three different experiments, stacks of 97 images were segmented by applying the classifier. A FIJI macro was generated to count IP-positive cells, involving successive steps of processing, including conversion to a mask, watershed segmentation, and analysis of particles. The number of IP-positive cells for each image was then determined and associated with specific time points to analyze cell death over time. From these data, we determined DCT50 by obtaining a percentage from the number of dead cells, then the time corresponding to 50% dead cells was calculated using R Software.

## Supporting information

All supplemental tables and figures

## Acknowledgement

The authors thank the Platform of Cell and Tissue Imaging (PICT) for imaging core facilities. We would like to deeply thank Dr Christophe Audebert and Dr Delphine Beury for their advice and technical assistance.

