## Supplementary material for "The *panC*-encoded pantothenate synthetase to tackle carbapenem-resistant OprD *Pseudomonas aeruginosa* mutant revealed through Tn-Seq": All supplemental tables and figures

### Supplementary data

**Table S1: Bacterial strain used in this study**

| Strain | Genotype and characteristics | Reference |
|---|---|---|
| ***P. aeruginosa*** | | |
| PA14 WT | Reference strain (*P. aeruginosa* PA14 UCBPP) | L.G. Rahme (54) |
| PA14Δ*oprD* | *P. aeruginosa* PA14 UCBPP with clean deletion of the *oprD* gene | D. Skurnik (11,12) |
| PA14/pRS2022005 (pEXG2Δ*fahA*) | Strain with a pEXG2 plasmid containing the Up and Down sequences of *fahA* gene for deletion in PA14 | This work |
| PA14Δ*oprD*/pRS2022005 (pEXG2Δ*fahA*) | Strain with a pEXG2 plasmid containing the Up and Down sequences of *fahA* gene for deletion in PA14 | This work |
| PA14Δ*fahA* | *P. aeruginosa* PA14 UCBPP with clean deletion of the *fahA* gene | This work |
| PA14Δ*oprD*Δ*fahA* | *P. aeruginosa* PA14 UCBPP with clean deletion of the *oprD* gene and *fahA* gene | This work |
| PA14/pRS2022006 (pEXG2Δ*panC*) | Strain with a pEXG2 plasmid containing the Up and Down sequences of *panC* gene for deletion in PA14 | This work |
| PA14Δ*oprD*/pRS2022006 (pEXG2Δ*panC*) | Strain with a pEXG2 plasmid containing the Up and Down sequences of *panC* gene for deletion in PA14 | This work |
| PA14 WT/ pJMP1237::sg_CTL | CRISPRi module with sgRNA with non-targeting spacer in plasmid pJMP1237 and integrated into the PA14 chromosome at att Tn7 after the *glmS* gene (6.516.367-6.518.202). | This work |
| PA14 *ΔoprD*/ pJMP1237::sg_CTL | CRISPRi module with sgRNA with non-targeting spacer in plasmid pJMP1237 and integrated into the PA14 chromosome at att Tn7 after the *glmS* gene (6.516.367-6.518.202). | This work |
| PA14 WT/ pJMP1237::sg_*panC* | CRISPRi module with sgRNA targeting the *panC* gene constructed in the plasmid pJMP1237 and integrated into the PA14 chromosome at att Tn7 after the *glmS* gene (6.516.367-6.518.202). | This work |
| PA14*ΔoprD*/ pJMP1237::sg_*panC* | CRISPRi module with sgRNA targeting the *panC* gene constructed in the plasmid pJMP1237 and integrated into the PA14 chromosome at att Tn7 after the *glmS* gene (6.516.367-6.518.202). | This work |
| PA14 WT /pJMP1237::sg_*fahA* | CRISPRi module with sgRNA targeting the *fahA* gene constructed in the plasmid pJMP1237 and integrated into the PA14 chromosome at att Tn7 after the *glmS* gene (6.516.367-6.518.202). | This work |
| PA14Δ*oprD* /pJMP1237::sg_*fahA* | CRISPRi module with sgRNA targeting the *fahA* gene constructed in the plasmid pJMP1237 and integrated into the | This work |

| | | |
|---|---|---|
| | PA14 chromosome at att Tn7 after the *glmS* gene (6.516.367-6.518.202). | |
| ***E. coli*** | | |
| E. coli DH5α/pEXG2 | Strain with the native plasmid pEXG2 | |
| *E. coli* DH5α/pRS2022005 (pEXG2Δ*fahA*) | Strain with a pEXG2 plasmid containing the Up and Down sequences of *fahA* gene for deletion in PA14 | This work |
| DH5α/pRS2022006 (pEXG2Δ*panC*) | Strain with a pEXG2 plasmid containing the Up and Down sequences of *panC* gene for deletion in PA14 | This work |
| *E. coli* MFD DAP-*pir*+/pJMP1237 | Strain used as donor for tri-parental mating with the plasmid containing the CRISPRi module. | This work |
| MFD DAP- pir+ /pJMP1039 | Strain used as donor for tri-parental mating with the plasmid containing transposase. | This work |

**Table S2: Quality metrics obtained with the Transit stats function from WIG files generated with bowtie mapper**

| | S1_RUN1 | S3_RUN1 | S1 | S2 | S5 | S6 |
|---|---|---|---|---|---|---|
| **Number of reads** | 8 003,192 | 8 713,300 | 6 806,023 | 10 967,540 | 9 955,879 | 1 148,038 |
| **Mapped_read** | 6 714,836 | 7 269,135 | 5 059,919 | 8 860,617 | 7 197,141 | 786,779 |
| **%mapped_reads** | 83.90 | 83.43 | 74.34 | 80.9 | 72.29 | 68.53 |
| **Density** | 0.599 | 0.661 | 0.643 | 0.659 | 0.674 | 0.539 |
| **mean_CT** | 66.4 | 71.9 | 50.0 | 87.6 | 71.2 | 7.8 |
| **NZ_mean** | 110.9 | 108.8 | 77.8 | 132.9 | 105.7 | 14.4 |
| **NZmedian** | 56 | 51 | 35 | 55 | 47 | 6 |
| **Max_ct** | 3284 | 3942 | 10661 | 17109 | 4579 | 964 |
| **Total_cts** | 6 704,929 | 7 259,889 | 5 049,342 | 8 841,568 | 7 187,530 | 785,172 |
| **Skewness** | 4.1 | 4.4 | 14.7 | 11.9 | 4.9 | 6.6 |
| **Kurtosis** | 30.4 | 36.9 | 857.8 | 590.6 | 46.6 | 98.8 |
| **Pickands_tail_index** | 0.155 | 0.133 | 0.182 | 0.237 | 0.245 | 0.271 |

**Table S3: Quality metrics obtained with the Transit stats function from WIG files generated with BWA**

| | S1_RUN1 | S3_RUN1 | S1 | S2 | S5 | S6 |
|---|---|---|---|---|---|---|
| **#reads** | 8003192 | 8713300 | 6806023 | 10967540 | 9955879 | 1148038 |
| **#trimmed_reads** | 8002396 | 8712179 | 6805509 | 10966681 | 9954917 | 1147948 |
| **#mapped_reads** | 7573647 | 8100871 | 5927151 | 10068452 | 8444399 | 938621 |
| **%reads_mapped** | 94.63 | 92.97 | 87.09 | 91.80 | 84.82 | 81.76 |
| **density** | 0.660 | 0.720 | 0.693 | 0.719 | 0.730 | 0.571 |
| **mean_ct** | 73.5 | 78.5 | 55.9 | 96.3 | 78.4 | 8.7 |
| **NZmean** | 111.3 | 109.0 | 80.8 | 133.9 | 107.5 | 15.3 |
| **NZmedian** | 53 | 48 | 34 | 52 | 45 | 6 |
| **max_ct** | 235553.0 | 160437.0 | 185711.0 | 251729.0 | 115348.0 | 20780.0 |
| **total_cts** | 7420002 | 7919795 | 5645965 | 9717220 | 7916904 | 880388 |
| **skewness** | 246.9 | 240.5 | 248.0 | 246.4 | 194.8 | 200.6 |
| **kurtosis** | 62785.3 | 62356.3 | 64124.8 | 64368.2 | 46652.0 | 45145.6 |
| **pickands_tail_index** | 0.173 | 0.186 | 0.630 | 0.551 | 0.537 | 0.605 |

**Table S4: Summary of the studies on essential genes in *P. aeruginosa***

| Dataset | Year | #EG | Strain | Media | Ref |
|---|---|---|---|---|---|
| **Liberati** | 2006 | 335 | PA14 | LB | (19) |
| **Skurnik** | 2013 | 636 | PA14 | LB | (11) |
| **Turner** | 2015 | 314 | PA14 | BHI Agar | (20) |
| **Lee** | 2015 | 314 | PAO1 | LB | (21) |
| **Poulsen** | 2019 | | PA14 | LB | (65) |
| *FWER* | | 437 | | | |
| *FDR* | | 596 | | | |
| *TRANSIT* | | 316 | | | |
| *CORE* | | 321 | Core pangenome *P. aeruginosa* | | |
| *Small* | | 93 | Intersection of 5 pangenomes studies | | |

**Table S5: Plasmids used in this study**

| Plasmids | Characteristics | Reference |
|---|---|---|
| **Gene deletion** | | |
| **pEXG2** | Allelic exchange vector with pBR origin, gentamicin reistance, *sacB* | Arne Rietsch (60) |
| **pEXG2::Δ*fahA*** | pEXG2 containing the Up and Down sequences (500bp) of *fahA* gene of PA14 for deletion | This work |
| **pEXG2::Δ*panC*** | pEXG2 containing the Up and Down sequences (500bp) of *panC* gene of PA14 for deletion | This work |
| **CRISPRi/dCas9** | | |
| **pJMP1039** | Plasmid containing the transposase allowing the integration of the CRISPRi module into the PA14 genome | Gift from Carol Gross & Jason Peters & Oren Rosenberg |
| **pJMP1237** | Plasmid containing the CRISPRi module framed by Tn7 restriction sites. Gentamicin resistance gene associated with the CRISPRi module. | Gift from Carol Gross & Jason Peters & Oren Rosenberg |
| **pJMP1237::sg_CTL** | pJMP1237 containing the 20bp non targeting spacer in PA14 to use as control during inhibition conditions | This work |
| **pJMP1237::sg_*panC*** | pJMP1237 containing the 20bp spacer of the PA14 *panC* gene for inhibition of the gene by CRISPRi | This work |
| **pJMP1237::sg_*fahA*** | pJMP1237 containing the 20bp spacer of the PA14 *fahA* gene for inhibition of the gene by CRISPRi | This work |

**Table S6: List of the primers used for RT-qPCR**

| Primer | Sequence | Use |
|---|---|---|
| **DYH115** | 5' CAT-AAT-ATC-TCA-TTT-CAC-TAA-ATA-ATA-GTG-AAC-GGC-AGG-TAA-GC 3' | Use to check fragments in pEXG2-designed by DYH |
| **DYH116** | 5' CAT-TCT-GCT-AAC-CAG-TAA-GGC-AAC-CCC-G 3' | To check fragments in pEXG2-designed by DYH |
| **PA14_pEXG2_F** | 5' AGC-TCG-AAT-TCG-GTA-CCT-TAA-TTA-ATT-TC 3' | Primers for gene deletion with PEXG2 by Gibson Assembly |
| **PA14_pEXG2_R** | 5' CCG-GGG-ATC-CTC-TAG-AGT-C 3' | Primers for gene deletion with PEXG2 by Gibson Assembly |
| ***fahA* up_fwd** | 5' CGA-CTC-TAG-AGG-ATC-CCC-GGG-AGC-TTG-ACC-ACT-CGC-CG 3' | Primer for *fahA* deletion in PA14 by Gibson Assembly, overlaping PA14_pEXG2_R |
| ***fahA* up_rev** | 5' AGA-GCT-TCA-TGG-GGT-TAT-CTC-CGT-TGC-G 3' | Primer for *fahA* deletion in PA14 by Gibson Assembly, overlaping *fahA*_down |
| ***fahA* down_fwd** | 5' AGA-TAA-CCC-CAT-GAA-GCT-CTA-TAC-CTA-CTA-CCG-TTC-CAC-CTC 3' | Primer for *fahA* deletion in PA14 by Gibson Assembly, overlaping *fahA*_up |
| ***fahA* down_rev** | 5' TAA-GGT-ACC-GAA-TTC-GAG-CTG-TGG-CCA-GGC-GTC-CAG-CG 3' | Primer for *fahA* deletion in PA14 by Gibson Assembly, overlaping PA14_pEXG2_F |
| ***panC* up_fwd** | 5' CGA-CTC-TAG-AGG-ATC-CCC-GGC-CTG-GCC-TTC-GAA-TCC-AG 3' | Primer for *panC* deletion in PA14 by Gibson Assembly, overlaping PA14_pEXG2_R |
| ***panC* up_rev** | 5' TCG-GCA-TGC-GTC-ATG-CGT-TGA-ATC-CGT-G 3' | Primer for *panC* deletion in PA14 by Gibson Assembly, overlaping *panC*_down |
| ***panC* down_fwd** | 5' CAA-CGC-ATG-ACG-CAT-GCC-GAG-CCC-CCG-G 3' | Primer for *panC* deletion in PA14 by Gibson Assembly, overlaping *panC*_up |
| ***panC* down_rev** | 5' TAA-GGT-ACC-GAA-TTC-GAG-CTG-TTG-GAA-TTC-GAT-GGC-GCA-CTT-ATT-TGA-ACA-TGA-AAA-AGC-C 3' | Primer for *panC* deletion in PA14 by Gibson Assembly, overlaping PA14_pEXG2_F |
| ***fahA*gene_F** | 5' CTG-TCT-GCT-CAA-CGA-CTG 3' | Primers for verification of the deletion of *fahA*gene |
| ***fahA*gene_R** | 5' CGT-TGT-CGT-CGT-AAA-GAT-AC 3' | Primers for verification of the deletion of *fahA*gene |
| ***panC*gene_F** | 5' AGA-CCG-TCC-GTG-AAC-TG 3' | Primers for verification of the deletion of *panC* gene |
| ***panC*gene_R** | 5' GGT-ATG-TGT-CGA-GGT-CTT-C 3' | Primers for verification of the deletion of *panC*gene |
| ***sigX_F*** | 5' AAC-CAG-AAT-CTC-CCG-ATC-TA 3' | Primers for verification of the deletion of *fahA* gene |

| | | |
|---|---|---|
| ***sigX_R*** | 5' GCA-AGT-CGA-AGT-TCA-AAA-CA 3' | Primers for verification of the deletion of *fahA* gene |
| ***algU_F*** | 5' GGT-ATT-CAA-CTG-GGT-GAA-CT 3' | Primers for verification of the deletion of *panC* gene |
| ***algU_R*** | 5' CAA-GTT-CGA-TCC-TTC-CCA-A 3' | Primers for verification of the deletion of *panC* gene |
| ***cmpX_F*** | 5' GAC-GTA-GAT-TCC-TGC-AAT-GA 3' | Primer for RT- PCR of *sigX* gene |
| ***cmpX_R*** | 5' GGT-CAT-CAT-CAT-CAG-CAT-CT 3' | Primer for RT- PCR of *sigX* gene |
| ***cmaX_F*** | 5' AGG-AAA-AAG-CCG-GTG-ATT-AT 3' | Primer for RT- PCR of *cmpX* gene |
| ***cmaX_R*** | 5' GGG-ATA-TCT-ACT-CGC-AAC-TG 3' | Primer for RT- PCR of *cmpX* gene |
| ***fabD_F*** | 5' CAG-GTA-GTG-ATC-GCC-GGT 3' | Primer for RT- PCR of *fabD* gene |
| ***fabD_R*** | 5' CAG-CGT-ATC-GAG-ATC-AGC 3' | Primer for RT- PCR of *fabD* gene |
| ***fabG_F*** | 5' CCA-ACC-TGA-ACA-GTC-TCT-AC 3' | Primer for RT- PCR of *fabG* gene |
| ***fabG_R*** | 5' ATT-CAC-GGT-AAT-GGC-ACG 3' | Primer for RT- PCR of *fabG* gene |
| ***fabY_F*** | 5' GAA-GGG-TAT-CGA-CGT-CAT-C 3' | Primer for RT- PCR of *fabY* gene |
| ***fabY_R*** | 5' TCC-ATC-AGC-ACT-ACG-TAC-T 3' | Primer for RT- PCR of *fabY* gene |
| ***fabZ_F*** | 5' CTC-ATC-GCT-ATC-CTT-TCC-TG 3' | Primer for RT- PCR of *fabZ* gene |
| ***fabZ_R*** | 5' GCA-TCT-TGA-AAC-CGA-GGA-TA 3' | Primer for RT- PCR of *fabZ* gene |
| ***gyrB_F*** | 5’ AGC-TAC-GTC-ACC-TTC-AAC-CG 3’ | Primer for RT- PCR of *gyrB* gene |
| ***gyrB_R*** | 5’ ACA-GCT-GCT-CAG-GGT-TCA-TC 3’ | Primer for RT- PCR of *gyrB* gene |

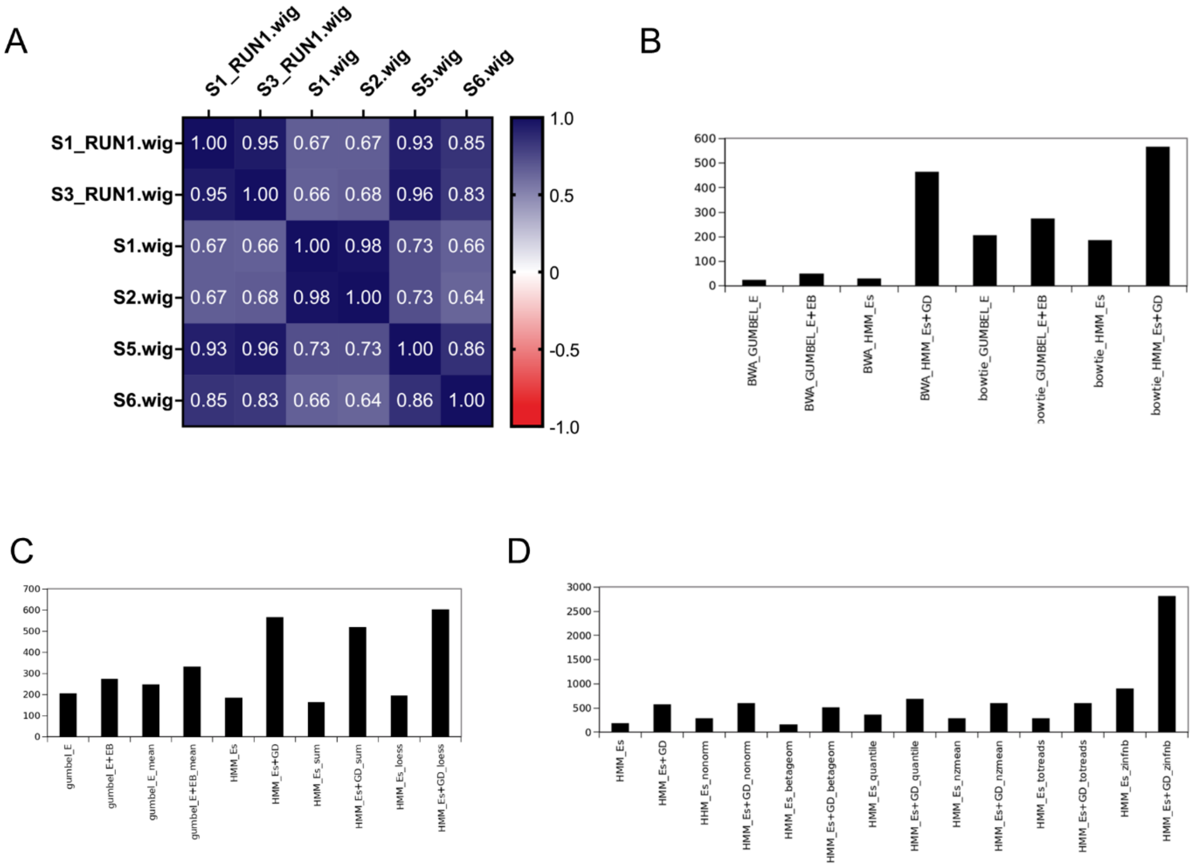


**Figure S1**: **Assessment of methods for TnSeq data analysis. (A)** Correlation matrix (Pearson coefficients) between the six samples used in TnSeq. Data were obtained with the corrplot R package from WIG file, plotted with GraphPad Prism. Dark blue represents highest correlation between two samples. **(B)** Impact of the mapper on EGs determination represented by the number of EGs identified with BWA and bowtie. **(C)** Impact of software parameters on EGs determination represented by the number of EGs identified with BWA and bowtie. **(D)** Impact of normalization on EGs determination represented by the number of EGs identified with BWA and bowtie. EG = Essential class, EB= Binomial class and GD = Growth Defect class.

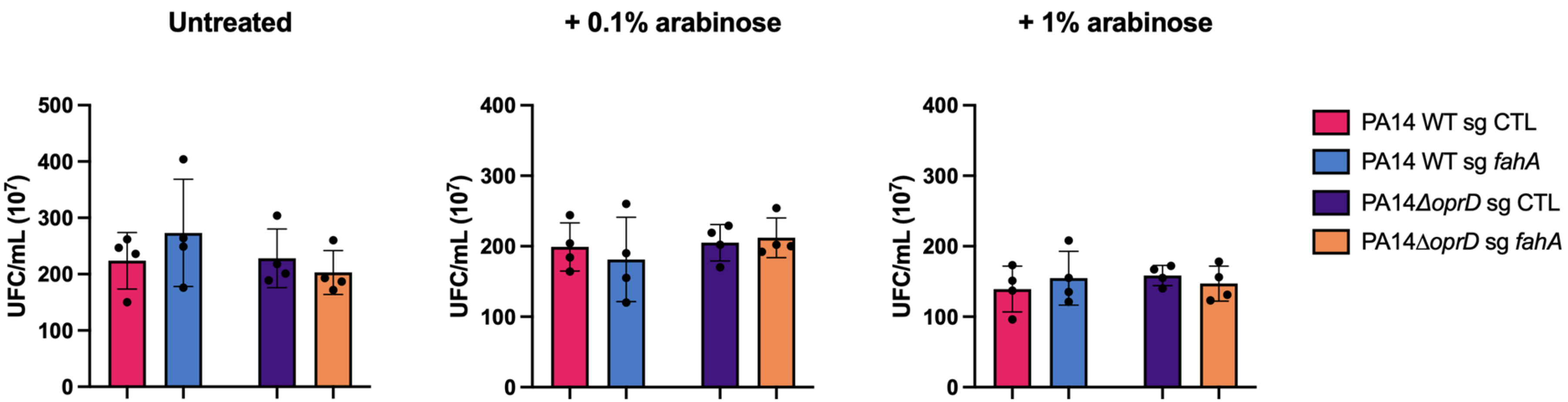


**Figure S2**: **Biological invalidation of *fahA* essentiality for PA14Δ*oprD***. Colony count of the PA14 CRISPRi stains after culture in LB media supplemented with various concentration of arabinose (0.1% or 1%) for 6 hours (n=4). Statistics were achieved by Mann-Whitney test.

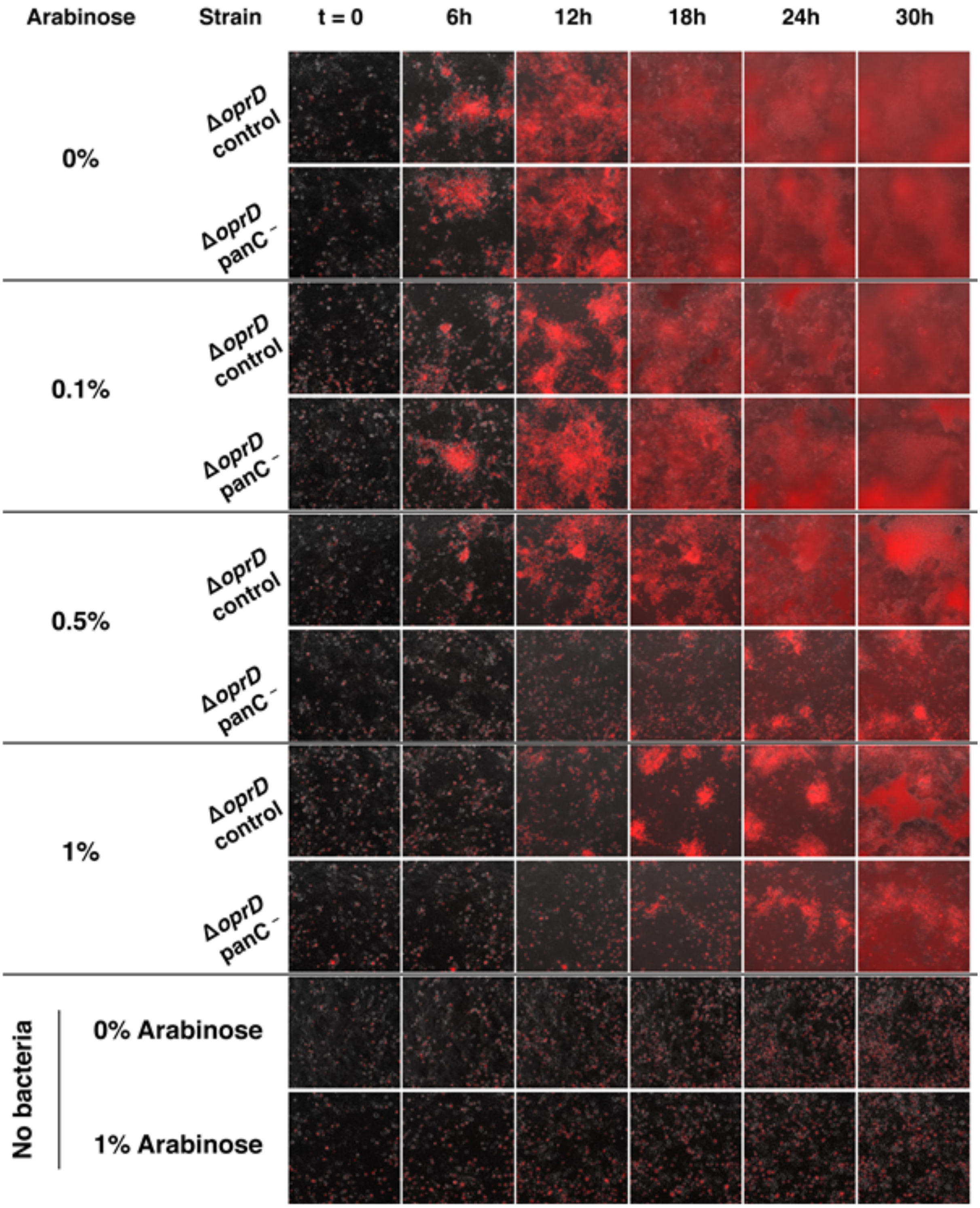


**Figure S3**: **16HBE cells dynamics of cell death following infection by CRISPRi PA14 strains.** Dynamics of cell death of 16HBE bronchial epithelial cells, infected by CRISPRi PA14 WT control, CRISPRi PA14 WT sg *panC*, CRISPRi PA14 Δ*oprD* control and CRISPRi PA14 Δ*oprD* sg *panC* strains. Images were taken using phase contrast microscopy and fluorescence microscopy (x20 objective). Cell death is visualized by the mortality marker propidium iodide (in red).
